# Performing Roux-N Y Gastric Bypass on Rabbits: Perspective Report to Evaluate Body Weight Changes Food Consumption, And the Malabsorption of Vitamin D3

**DOI:** 10.1101/2023.01.20.524848

**Authors:** Osaid Al Meanazel, Fars K. Alanazi, Mohammad Hailat, Wael Abu Dayyih, Ramadan Al-Shdefat, Mohammad Abu Assab, Riad Awad, Israa Al-Ani, Faiyaz Shakeel, Doaa H. Alshora, Mohamed A. Ibrahim

## Abstract

In 1998, Dr. Scopinaro published the first-ever known bariatric surgery, followed by reports by Buchwald and Oien in 2003 and 2013 [1,2]. Bariatric surgery (BS) is the most effective therapy against obesity, and recently it was recommended for type 2 diabetes as a therapeutic plan [3]. BS includes; Roux-en-Y gastric bypass (RYGB), sleeve gastrectomy, and laparoscopic adjustable gastric banding has gained the attention of most healthcare providers in elevation of the issue of obesity for its ease and fast results, especially with patients with mobility problems and those with high risk to develop chronic diseases [1,2]. Gastric bypass is considered the most common BS and a gold standard for weight loss, also known as Roux-en-Y gastric bypass (RYGB). The procedure of this BS is by creating a small pouch of the stomach (300 mL capacity) connected directly to the jejunum by bypassing the duodenum. RYGB has many advantages, such as; the alteration of guts hormones, which reduces appetite and hunger feeling, increases energy expenditure, causes significant long-term weight loss (60-80% of the excess body weight), decreases the amount of food consumed, and preserves 50% of the weight loss during this procedure. However, RYGB has some disadvantages. The most critical issue regarding this BS is altering the absorption of many medicines and nutrients, such as; vitamin D3 (Vit D3) [4,5]. (American Society for Metabolic and Bariatric Surgery. Bariatric Surgery Procedures. https://asmbs.org/patients/bariatric-surgery-procedures (accessed Feb 15, 2020). RYGB was performed on rodents before [6] but was never applied to rabbits. To determine the malabsorption procedure after the RYGB, the pharmacokinetics of the substance must be attained, and data must be calculated, such as; C_max_ and T_max_, to determine the absorption mutation. It was reported that pharmacokinetic studies on rodents could not be accurate due to the erratic absorption of the rodents [7]; thus, many studies have suggested using larger animals, such as rabbits, that do not address the same problem [8]. The current study investigates the ability to perform RYGB on rabbits and tests the effect of the surgery on body weight, food consumption, and the absorption of Vit D3.

## Animal choice and Ethical approval

The experimental animal care center kindly provided Male New Zealand rabbits, college of pharmacy, King Saud University, weighed 2.5-3.0 Kg and were individually housed in stainless steel cages under 12 h light cycle at room temperature 23 ± 3 oC. Standard rabbit food was purchased from the local markets and was available ad libitum unless otherwise mentioned. The Research Ethics Committee (REC) at King Saud University had approved all procedures regarding the animal surgery; approval references number KSU-SE-19-82.

## Pre-Operative Care

a. Food was restricted from the rabbits 12 hours before the surgery.
b. Anesthesia was applied through sevoflurane 5% via a tracheal tube (4.0 gauge).
c. An electric razor removed abdominal hair.
d. Rabbits were placed in a supine position after they had been anesthetized on an isothermal pad.
e. Before placing the rabbit’s snout in the nosecone, eye lubricant was applied to ensure no surgical complications.
f. Sevoflurane was maintained after the rabbits slept at 5.0% and O2 at 50%.
g. The skin of the abdomen was disinfected using a betadine solution.
h. Anesthesia was confirmed by pinching the rabbits between the toes of the hind leg.
i. Enrofloxacin, with a dose of 5.7 mg/Kg, was injected intraperitoneally as a surgery prophylaxis to avoid any post-operative infections.

**Figure 1.**
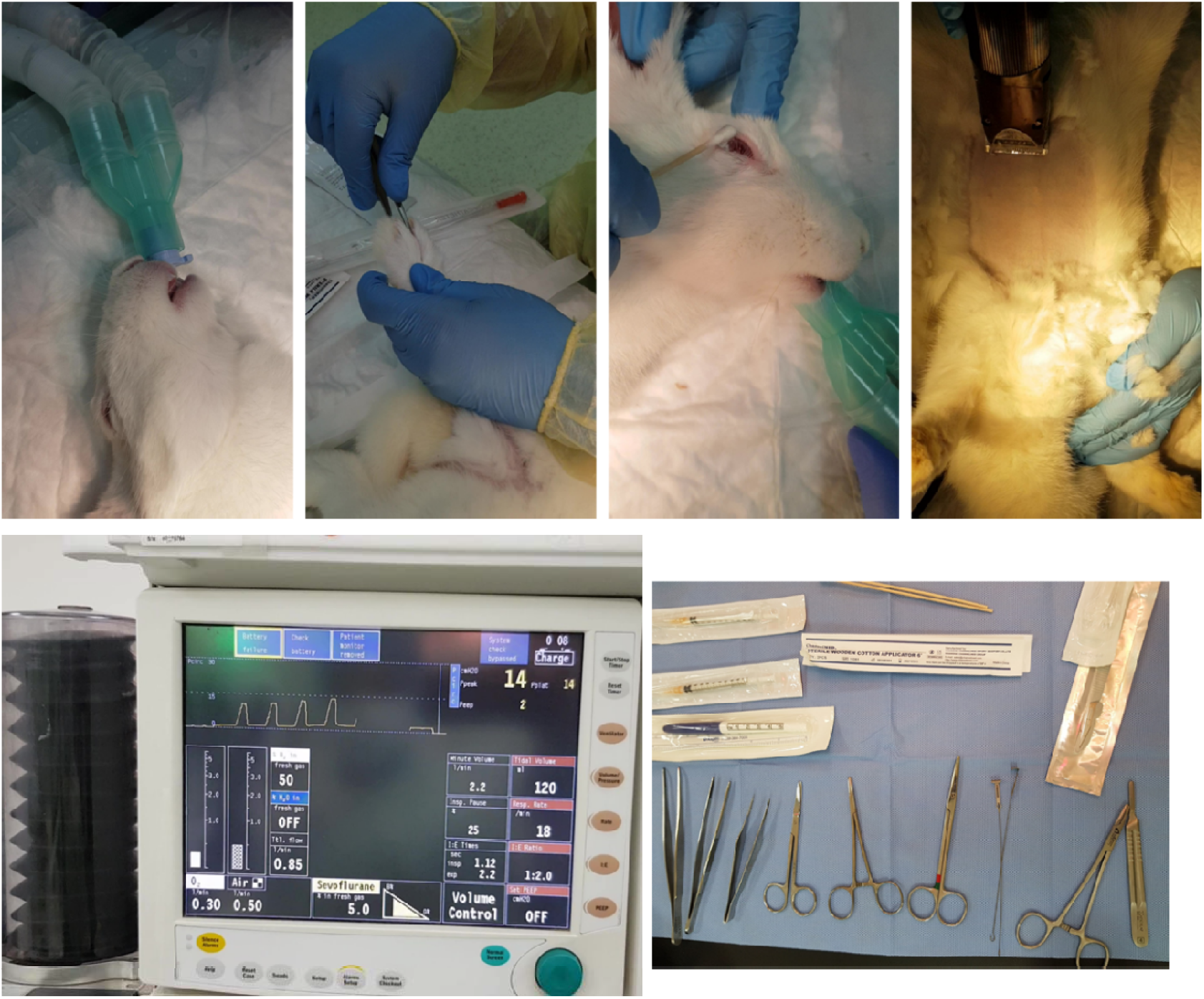
Pre-operative procedure.

## Median Laparotomy

a. The midline incision was performed using blade no. 10, 5 cm below the xiphoidal process.
b. Metzenbaum scissor was used to mobilize the abdominal skin from the abdominal muscles.
c. The midline incision was 10 cm long and included the abdominal muscles.

**Figure 2.**
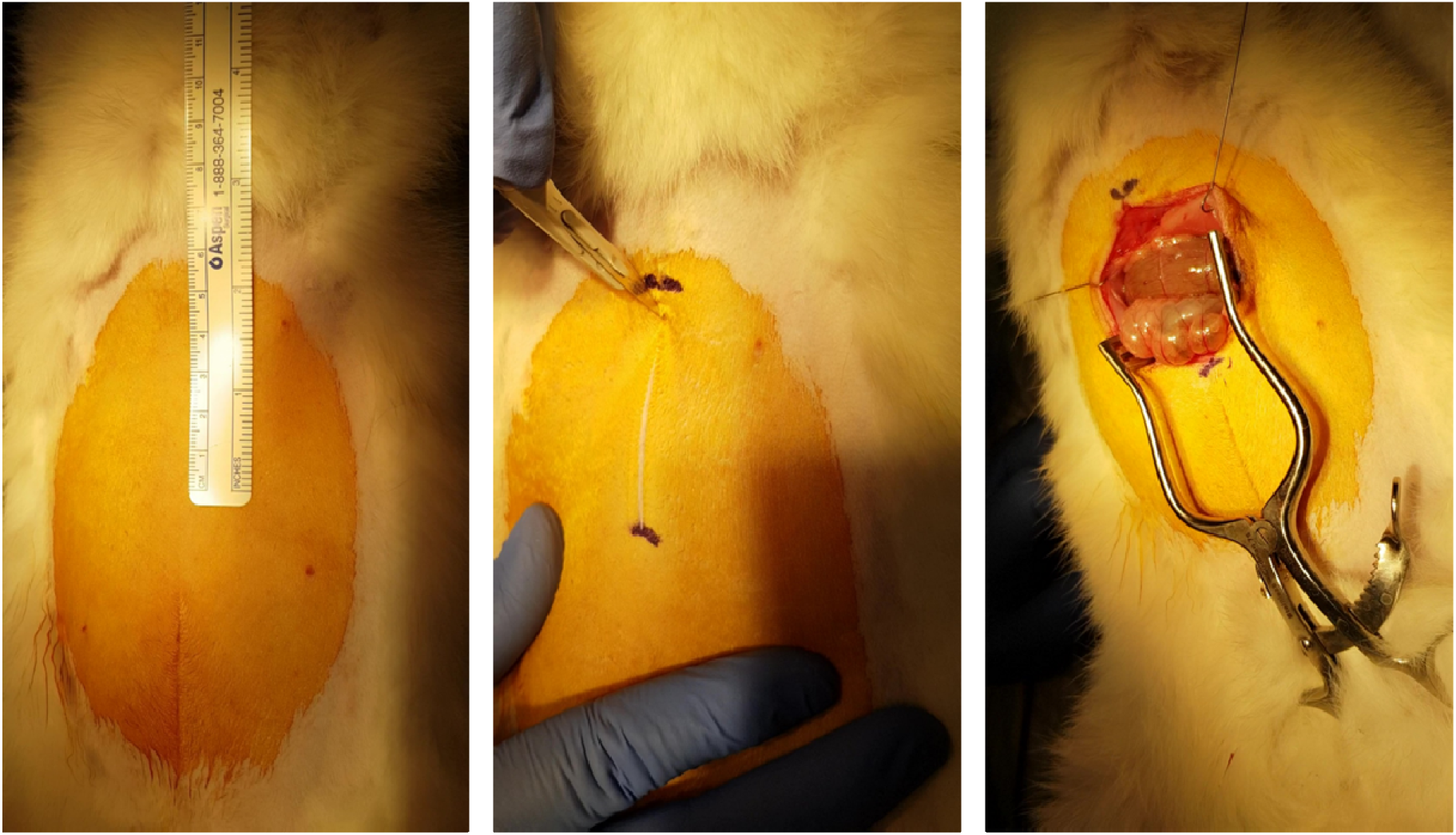
Retractors were installed on the bases of the best operation site vision.Median Laparotomy.

## Jejunostomy

a. The jejunum was identified under the colon, 10 cm away from the end of the duodenum, two points along the jejunum were restrained with hemostatic forceps and separated using a thermal cutter device.
b. The upper side of the jejunum was fused to the ileum, while the lower part was placed near the esophagus.

**Figure 3.**
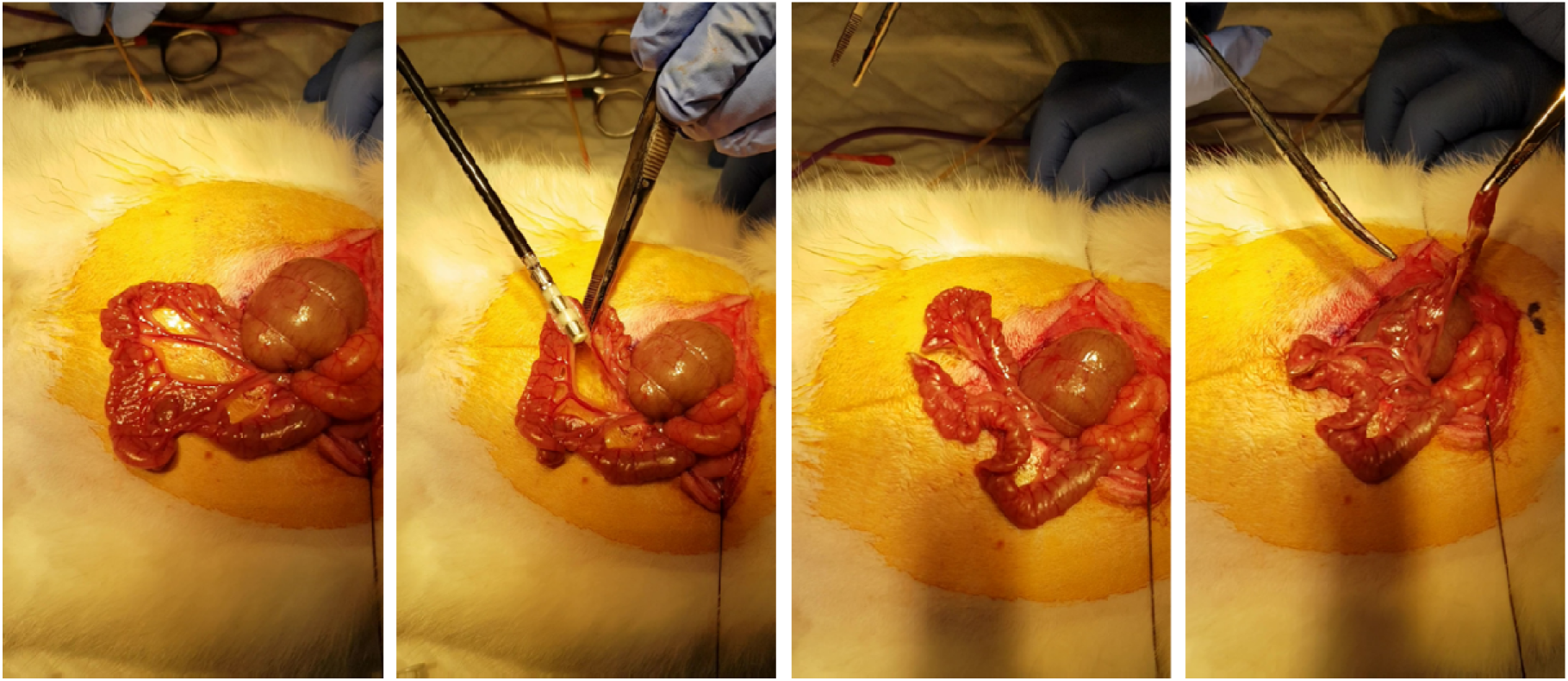
Jejunostomy.

## Gastric Pouch

a. At 2 cm from the end of the esophagus, the opening of the stomach, a double-layer suture was applied to prevent any leakage of the gastric content. The gastric pouch, along with the esophagus, was separated from the stomach; after that, the stomach was fixed at the internal wall of the abdomen with a suture to ensure the immobility of the internal organs due to the gab may produce by removing the stomach.

**Figure 4.**
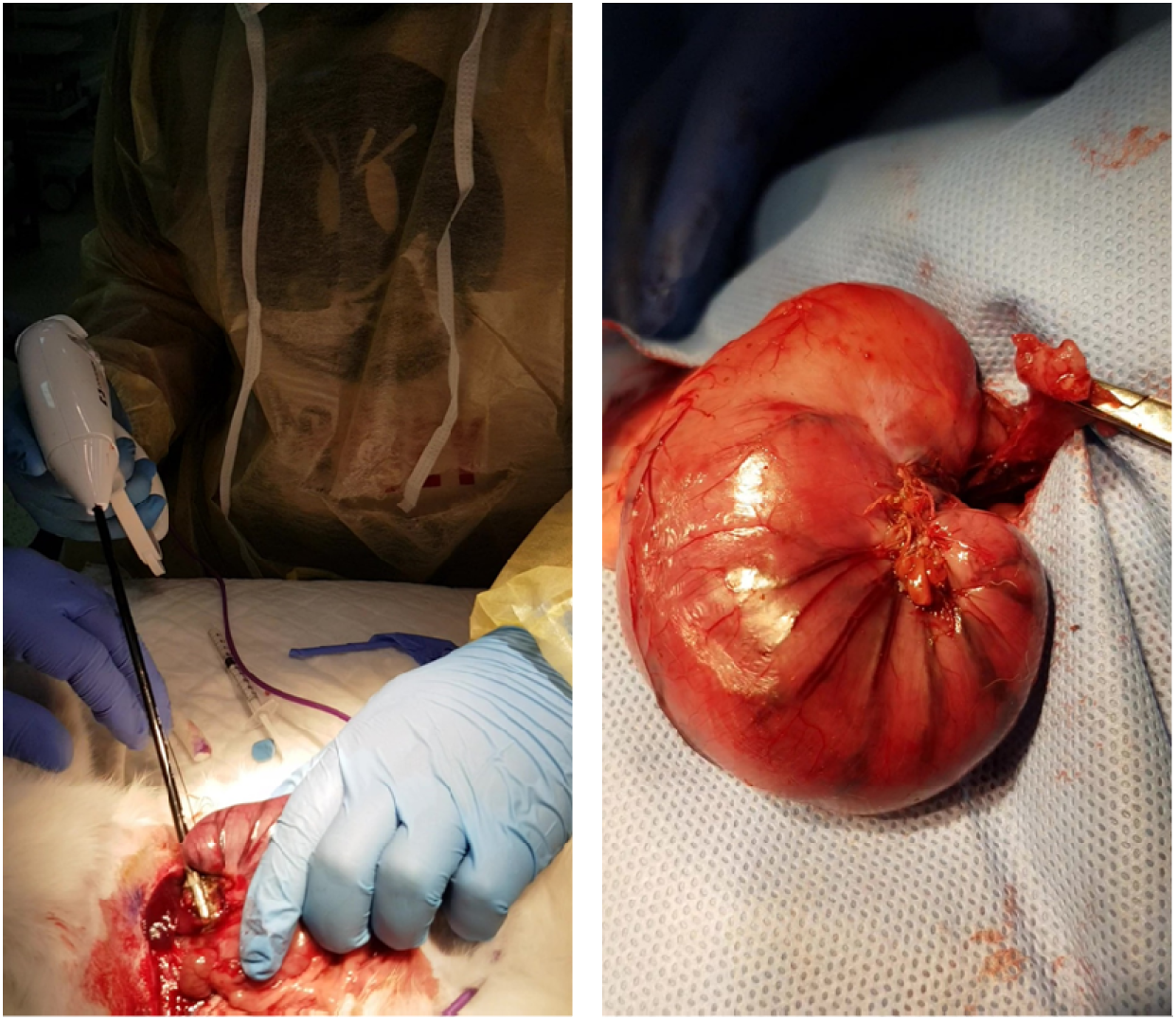
Gastric pouch.

## Gastro-Jejunostomy

a. The jejunum’s lower part was retrieved and positioned next to the gastric pouch.
b. Performing a gastro-jejunostomy by end-to-side anastomosis.
c. Suturing starts from back to front.

**Figure 5.**
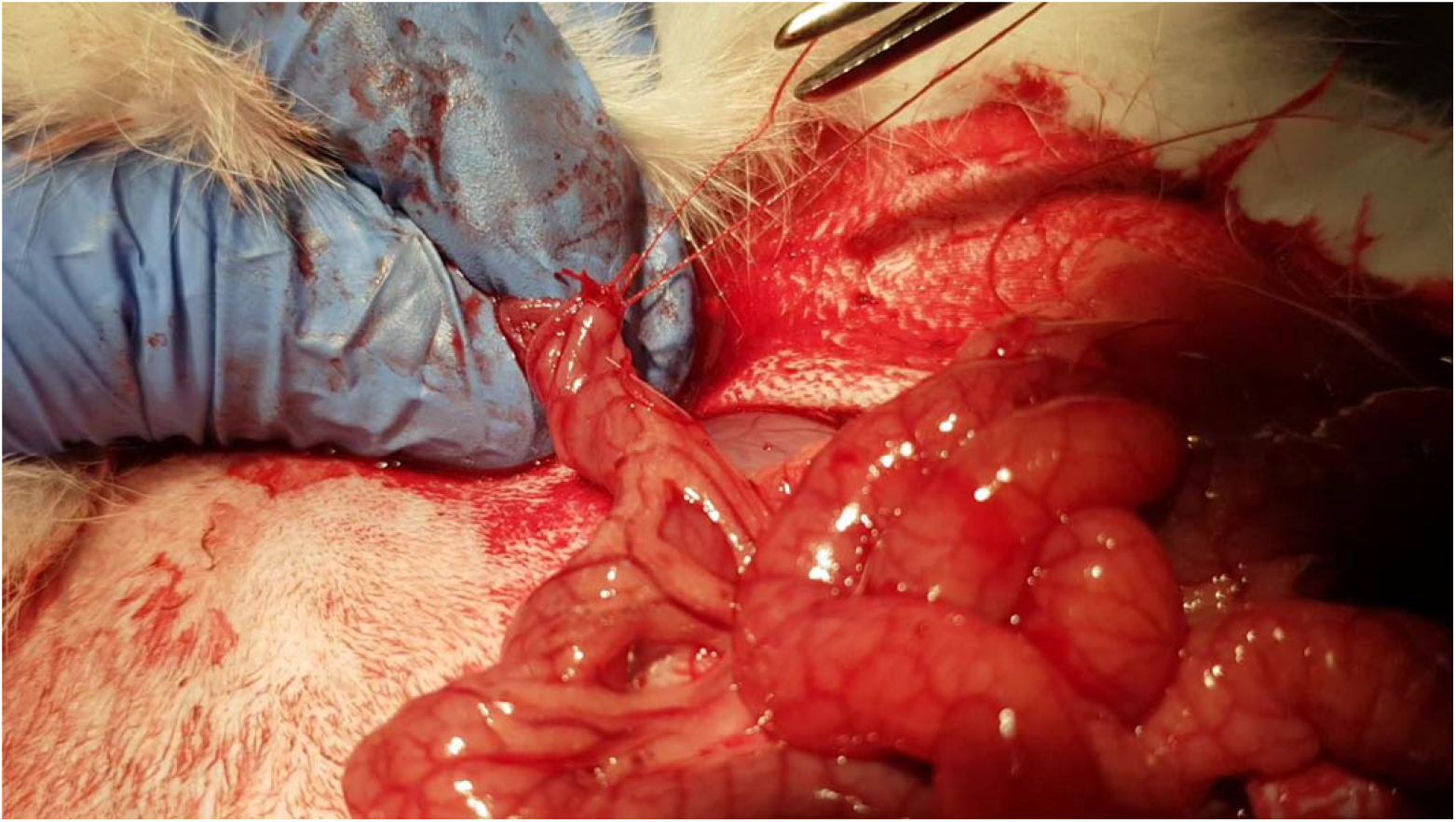
Gastro-Jejunostomy.

## Abdominal Closure

a. Sevoflurane was reduced to 2.5%
b. PDS 4-00 suture was used to close the abdominal muscle layer.
c. Sevoflurane was further reduced to 1.0%.

**Figure 6.**
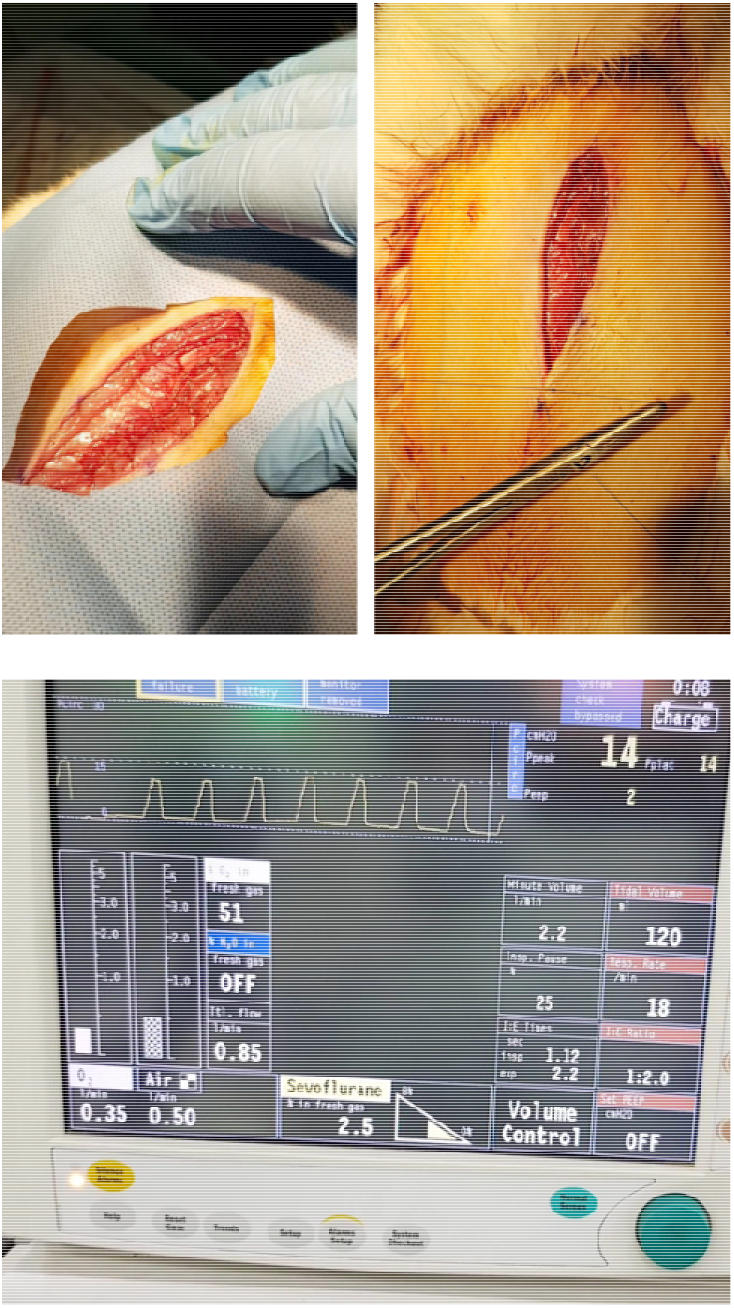
Vicryl 4-0 interrupted suture was utilized to close the surgery site skin.abdominal closure.

## Post-operative care

a. Sevoflurane was stopped while O2 continued.
b. The subcutaneous layer of the surgery site was injected with warm normal saline to avoid any dehydration
c. The rabbit was placed under a red light until it was fully recovered.
d. Returning the rabbits to their cage.
e. All rabbits that underwent the RYGB were subjected to fluid therapy for three days after the surgery.

**Figure 7.**
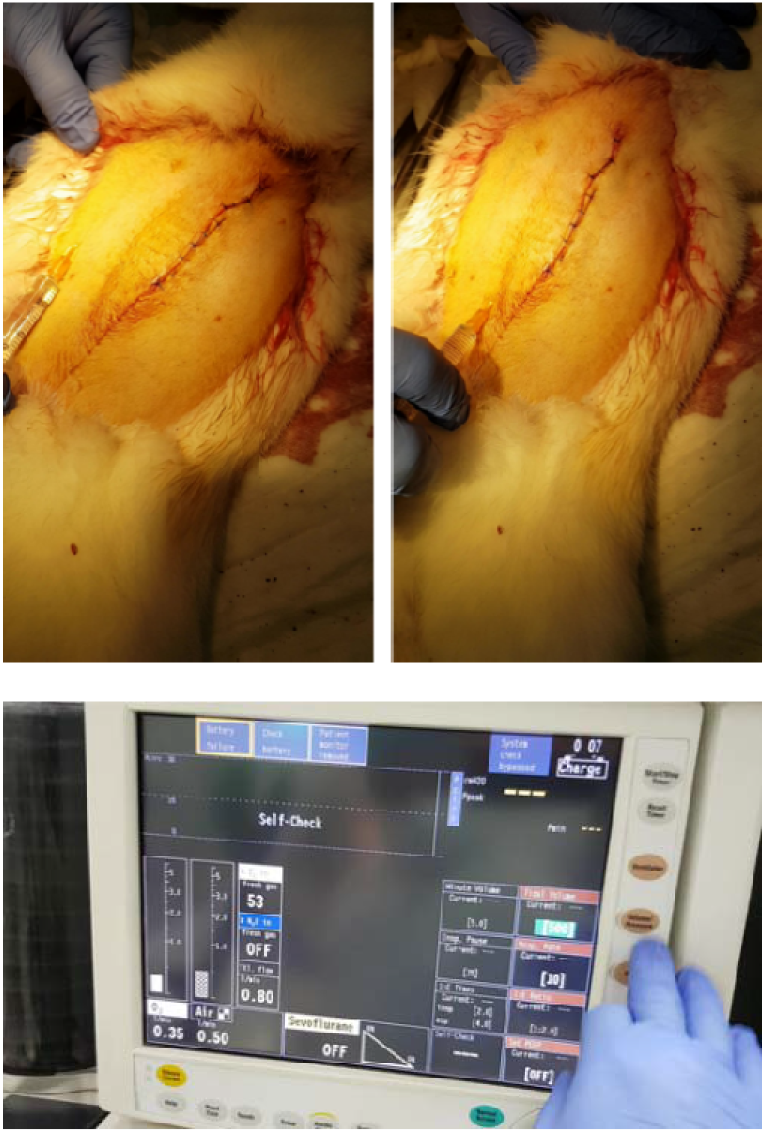
Post-operative procedure.

## Representative results

### Animals and Ethical Approval

The experimental animal care center kindly provided Male New Zealand rabbits, college of pharmacy, King Saud University, weighed 2.5-3.0 Kg and were individually housed in stainless steel cages under 12 h light cycle at room temperature 23 ± 3 °C. Standard rabbit food was purchased from the local markets and was available *ad libitum* unless otherwise mentioned. The Research Ethics Committee (REC) at King Saud University had approved all procedures regarding the animal surgery; approval references number KSU-SE-19-82.

### Body Weight

24 male New Zealand rabbits were used to evaluate the bodyweight change due to the RYGB. The rabbits were divided into two groups, G1 represents the healthy rabbits, and G2 symbolizes the RYGB rabbits. The rabbits’ weight was scaled before the surgery, after one week, two weeks, three weeks, and finally, four weeks. Fig. 32 represents the body weight changes during the period of the test. As for the G1 before the surgery (week 0), the average weight was 2.88 ± 0.22 Kg and for the G2 was 2.91 ± 0.28 Kg with no significant difference (P > 0.05). Nevertheless,after performing RYGB surgery on G2, the average body weight was significantly decreased to reach almost 2.17 ± 0.52 Kg for G2, which is considered a highly significant loss of body mass in contrast to the G1, 3.17 ± 0.18 Kg as shown in Fig. 32. this result came in agreement with previous studies that show the ability of RYGB weight loss and maintain the lost body mass [9–11].

**Figure 8.**
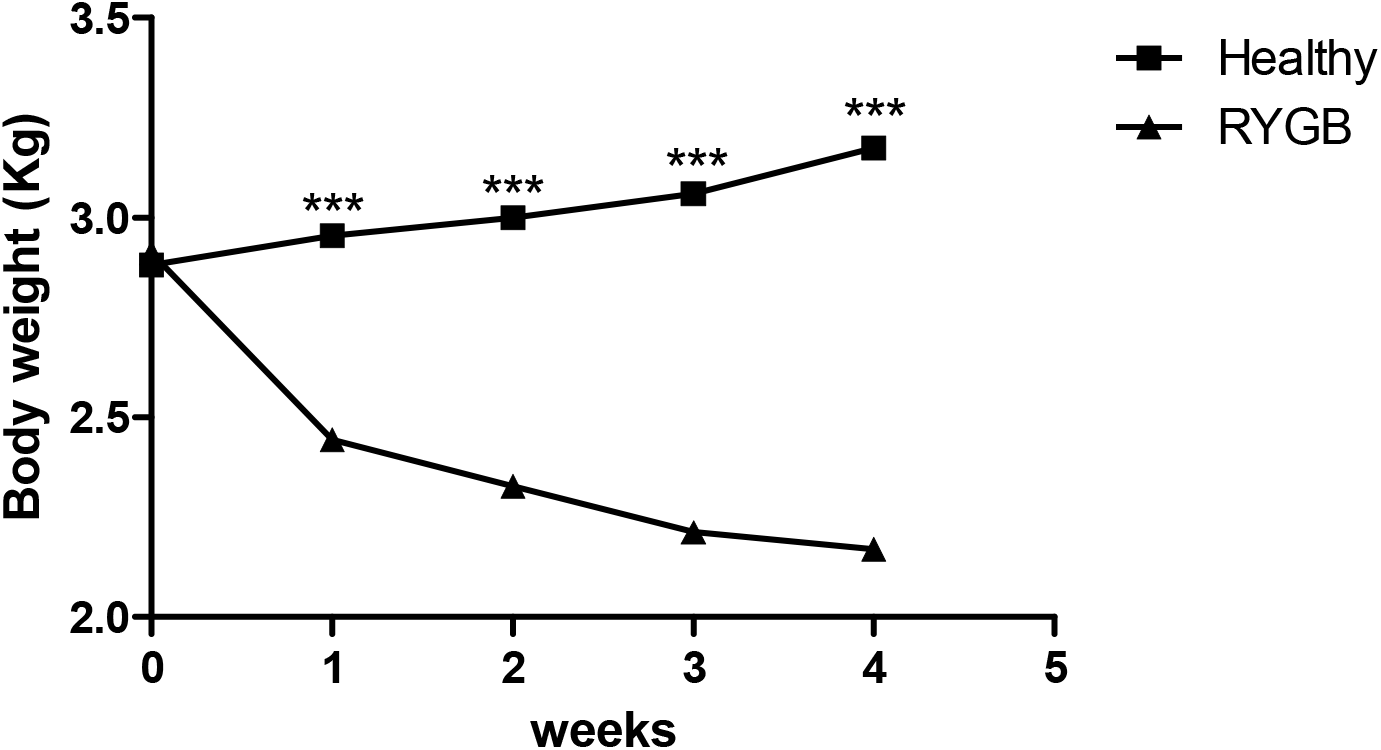
Bodyweight measurement for healthy and RYGB rabbits. (***) indicates a significant difference (P < 0.001).

### Food Consumption

Food consumption was evaluated throughout the study period (4 weeks). Each rabbit received 150 gm of food in the morning of each Sunday of the study period, and by the next day, morning food was collected, and the consumption was measured for individual rabbits to determine the average consumption of the whole group. Fig. 33 shows the average consumption (gm) of food for G1 and G2. It is obvious from G2 that the food consumption results came in parallel with the bodyweight results, as it indicates a consistent reduction, unlike the healthy rabbits (G1). The four weeks results exhibited a significantly lower food consumption than before the surgery (0 weeks) with a p-value of less than 0.001. On the other hand, G1 showed no significant difference between the week’s food consumption and thus excluded any other factors that may contribute to the food intake reduction other than RYGB. These results agreed with many studies that indicate the reduction of food intake after RYGB [10,12,13].

**Figure 9.**
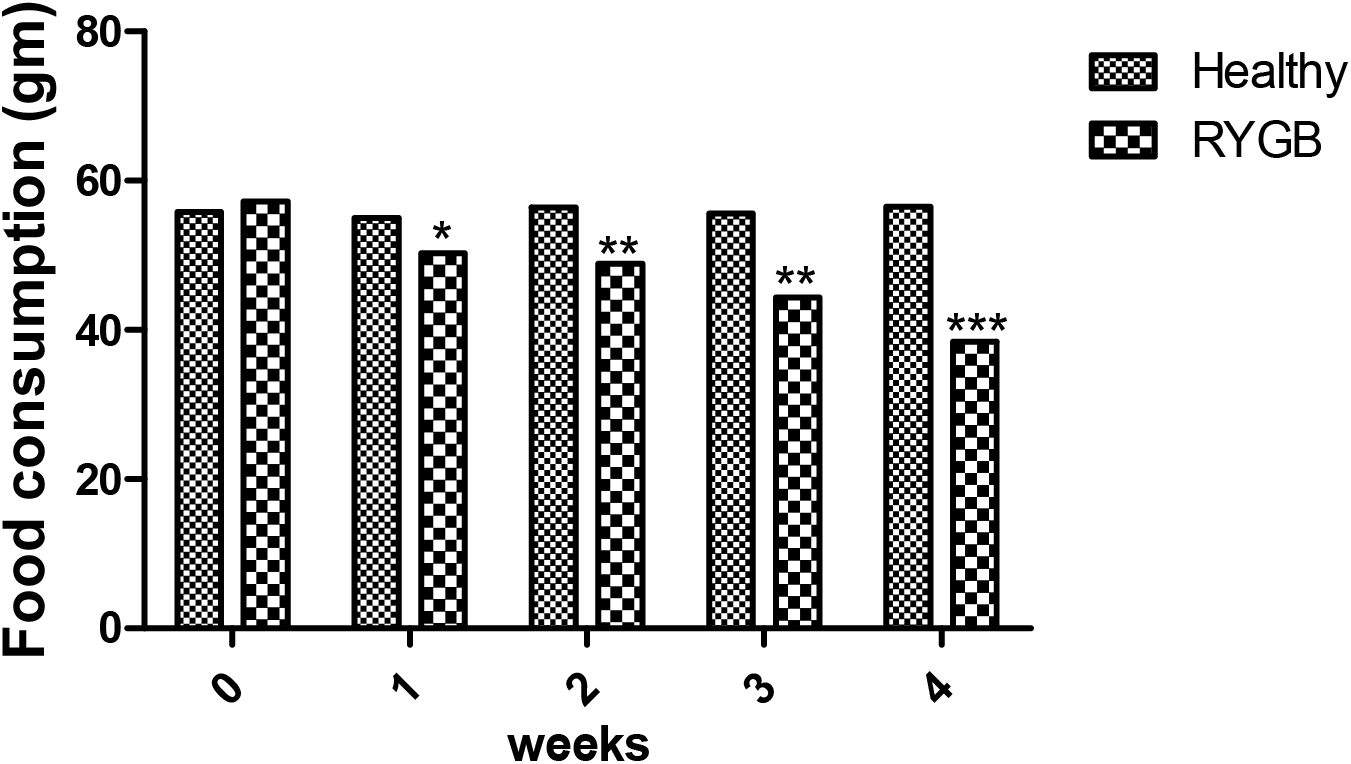
Food consumption per week for healthy and RYGB rabbits. (*) indicates a significant difference (P < 0.05), and (**) indicates a significantly different (P < 0.01). (***) indicates a significant difference (P < 0.001).

### Malabsorption Evaluation after RYGB

Although BS is a very efficient way of reducing weight, it affects the ability of drugs and nutrient absorption, leading to many cases of malnutrition (Ruiz-Tovar et al., 2013). One of the necessary elements that suffer from BS is vitamin D3 (Vit D3) (Ruiz-Tovar et al., 2013). Hence, many studies discussed decreased bone mineral density (BMD) after BS, mainly due to RYGB. Interestingly, most of them justified this consequence because of the malabsorptive procedure that occurs postoperatively [14,15].

To evaluate the malabsorption of Vit D3, two groups of Rabbits, G1(healthy) and G2 (RYGB), consisting of 3 rabbits, were administered orally with a commercial product of Vit D3. The dose of Vit D3 was calculated as the following equation:

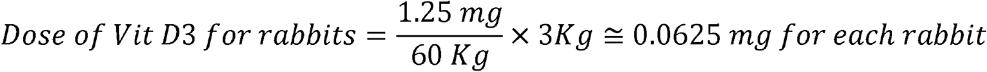

An amount equal to 0.0625 mg was suspended in 0.5% Carboxymethyl cellulose (CMC) and administered orally to each rabbit via oral gavage. Whole blood (amount ~ 500 ul) was collected through the ear vein using a winged infusion set (25 gauge) into lithium heparinized test tubes (3 ml) and was fixed for the whole period of the blood collection. Seven samples were withdrawn from each rabbit at times 0, 2, 4, 8, 14, 20, and 24 h. all blood samples were centrifuged at 6000 for 2 minutes to separate the plasma. The obtained plasma was kept in a −80 °C refrigerator for the pharmacokinetic analysis.

Samples were taken out of the refrigerator and kept at room temperature to melt and be ready to use. To analyze Vit D3 concentration in plasma, LIAISON® XL (Diasorin, Borsa Italiana S.p.A) chemiluminescence analyzer with LIAISON® 25 OH Vitamin D TOTAL Assay kit involved in it was utilized.

**Figure 10.**
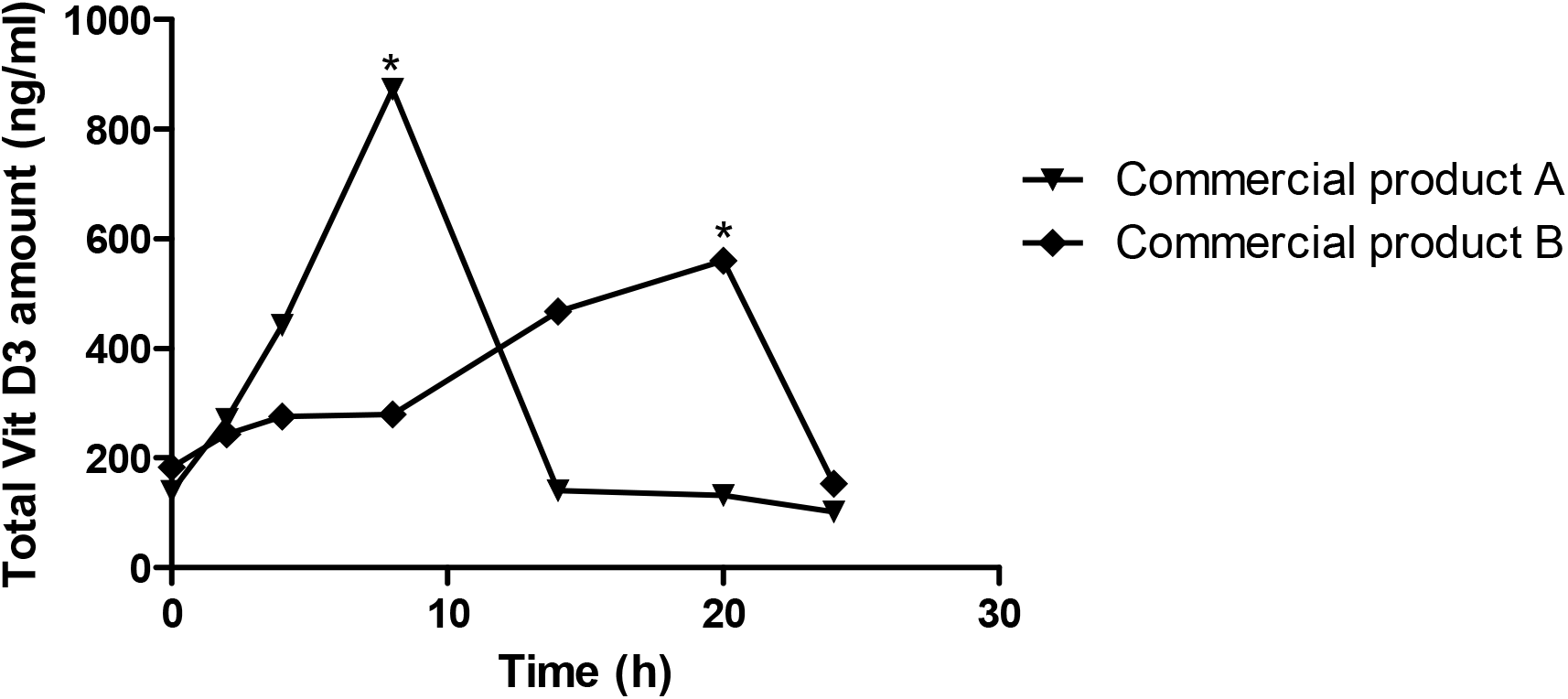
The Vit D3 plasma concentration curve for 24 h of commercial product in (A) healthy and (B) RYGB rabbit. (*) indicates a significant value (P < 0.05).

Fig.34 shows the total Vit D3 plasma concentration curves for 24 hs. The results show a significant delay of the T_max_ and a significant decrease in the C_max_ due to RYGB. T_max_ in the healthy rabbits (A) was eighth, while after the RYGB, it was delayed significantly (P < 0.05) to (B) 20 hs, with a decrease in Vit D3 C_max_ from 873.06 ± 218.8 ng/ml in the healthy animals, to 559.72 ± 224.81ng/ml after the RYGB. These results show the effect of RYGB as it proves the malabsorption of Vit D3 after RYGB.

## Conclusion

In conclusion, the study concluded the ability to develop an RYGB rabbit model, which also proved its efficacy for reducing weight; almost 30% reduction was obtained in the first four weeks of the surgery, and, more importantly, maintaining that mass loss. Food consumption was significantly reduced after the surgery. Moreover, the pharmacokinetic test dose proves the malabsorption of Vit D3, as it shows a significant delay in the T_max_, from 8 h in healthy to 20 h in the RYGB model (P < 0.05). In addition to the substantial decrease in C_max_, 873 ng/ml in healthy rabbits to almost 559 ng/ml in the RYGB model. This study encourages research to test the absorption changes for other substances and medications.

## Acknowledgments

We thank Prince Naif for Health Research Center and Prince Mutaib bin Abdullah Chair for Biomarkers of Osteoporosis, for their help in conducting the RYGB and analysis of the *in vivo* pharmacokinetic study.

